# Structural, functional and computational studies of membrane recognition by Plasmodium Perforin-Like Proteins 1 and 2

**DOI:** 10.1101/2021.12.16.473088

**Authors:** Sophie I. Williams, Xiulian Yu, Tao Ni, Robert J. C. Gilbert, Phillip J. Stansfeld

## Abstract

Perforin-like proteins (PLPs) play key roles in the mechanisms associated with parasitic disease caused by apicomplexans such as *Plasmodium* (malaria) and *Toxoplasma*. The *T. gondii* PLP1 (TgPLP1) mediates tachyzoite egress from cells, while the five *Plasmodium* PLPs carry out various roles in the life cycle of the parasite and with respect to the molecular basis of disease. Here we focus on *Plasmodium vivax* PLP1 and PLP2 (PvPLP1 and PvPLP2) compared to TgPLP1; PvPLP1 is important for invasion of mammalian hosts by the parasite and establishment of a chronic infection, PvPLP2 is important during the symptomatic blood stage of the parasite life cycle. Determination of the crystal structure of the membrane-binding APCβ domain of PvPLP1 reveals notable differences with that of TgPLP1, which are reflected in its inability to bind lipid bilayers in the way that TgPLP1 and PvPLP2 can be shown to. Molecular dynamics simulations combined with site-directed mutagenesis and functional assays allow a dissection of the binding interactions of TgPLP1 and PvPLP2 on lipid bilayers, and reveal a similar tropism for lipids found enriched in the inner leaflet of the mammalian plasma membrane. In addition to this shared mode of membrane binding PvPLP2 displays a secondary synergistic interaction side-on from its principal bilayer interface. This study underlines the substantial differences between the biophysical properties of the APCβ domains of Apicomplexan PLPs, which reflect their significant sequence diversity. Such differences will be important factors in determining the cell targeting and membrane-binding activity of the different proteins, in their different developmental roles within parasite life cycles.

## Introduction

*Plasmodium*, the parasitic causative agent of malaria, has a complex lifecycle involving two hosts, dominated by processes of migration through host tissue followed by cycles of intracellular replication, and *Plasmodium* Perforin-like proteins (*P*PLPs) have key roles in facilitating these processes [1–3]. PPLPs are found across apicomplexan species, and contain a conserved central membrane attack complex/perforin (MACPF) domain, as well as a unique C-terminal domain, named the Apicomplexan PLP C-terminal β-pleated sheet (APCβ) domain [4], and a variable N-terminal domain [5]. The MACPF superfamily represents an ancient homologous group of pore forming proteins, which function principally in cellular attack or defense. These proteins are secreted by a range of cells including bacterial and immune cells, bind to and oligomerize on cell membranes, and in this way form pores that typically result in target cell death [5–7]. Key members of this family are perforin-1 in cytotoxic lymphocytes [8, 9], bactericidal perforin-2 [10, 11], the complement system itself [12], and a range of toxins produced by organisms as disparate as stonefish [13] and the oyster mushroom [14]. Perforin family proteins may have a direct toxic effect (as complement does and also bacterial homologues) or be responsible for delivery of a secondary lethal agent (such as the delivery of granzymes by perforin-1) [9, 15].

The Apicomplexan PLPs (ApiPLPs), on the other hand, play important roles in cell traversal, invasion and egress by the parasites within their mammalian and insect hosts [1, 16–19]. The presence of MACPF domains in ApiPLPs suggests that membrane disruption by ApiPLPs is key to facilitation of these forms of cell behavior enabling infection and disease. *Plasmodium* species express 5 PLPs (PPLPs): two of them (PPLP1 and PPLP2) are involved in the mammalian host stage and the others (PPLP3-5) in the mosquito stage of the parasite life cycle. PPLP1 is implicated in cell traversal by sporozoites across the mammalian dermis and into the liver to invade hepatocytes and replicate within a parasitophorous vacuole (PV), as well as in the evasion of host phagocytic immune cells [1, 2, 3]. Knockout of PPLP1 significantly disrupts parasite transmission in *P. berghei* and *P. falciparum* [2, 20]. PPLP1 also plays a role in erythrocyte egress by the merozoite stage (gametocyte) of the *Plasmodium* life cycle, via membrane disruption [21]. PPLP2, too, has a function in gametocyte egress from erythrocytes for *P. berghei* [22] and *P. falciparum* [23]: exflagellation was disrupted in a PPLP2 knockout mutant, and was responsible for a subsequent inability to escape from erythrocytes. In contrast, PPLPs 3, 4 and 5 are all expressed during the parasite’s ookinete stage and facilitate traversal across the mosquito midgut epithelium [16, 17, 24–26]. Disruption of any of these three genes results in an inability by the ookinete to invade the mosquito midgut, suggesting distinct non-redundant roles for each of the three proteins [17, 27, 28]. They may thus act as part of a single complex in a similar sense to the assembly of the membrane attack complex [12].

Despite the significance of PPLPs in the malaria parasite life cycle, structural characterization has been lacking. A homologue from another member of the Apicomplexan family, *Toxoplasma gondii*, TgPLP1 facilitates parasite egress following intracellular replication [29]. X-ray crystallographic studies have previously revealed the structures of both the MACPF and APCβ domains of TgPLP1 [30, 31]. The TgPLP1 APCβ crystal structures revealed the basic characteristics of the domain, which consists of three repeats of a β-sandwich fold arranged in a prism architecture [30, 31]. It binds to target membranes via an extended loop at its base [30, 31] as similar features found in proteins such as perforin-1, perforin-2 and the CDCs do too [11, 32].

Here we present the first crystal structure of a *Plasmodium* PLP APCβ domain from *Plasmodium vivax* PLP1 at 3.15Å resolution, and confirm a shared domain architecture between TgPLP1 and the PvPLP APCβ domains. On the basis of these two structures we generate a homology model for the PvPLP2 APCβ domain and use molecular dynamics (MD) simulations and lipid binding assays to characterize their membrane tropism. Our results reveal that PvPLP1 does not bind to any membranes tested while PvPLP2 shows a binding preference for lipids of the inner plasma membrane leaflet similar to that of TgPLP1. Free energy calculations and *in silico* mutational studies then identify a common mechanism of membrane anchoring between TgPLP1 and PvPLP2 APCβ domains, clearly absent from PvPLP1. The variation in membrane interaction properties suggests possible functional and regulatory differences across the *Plasmodium* PLP family.

## Results

### Crystal Structure of the PvPLP1 C-terminal APCβ domain

The structure of PvPLP1 APCβ domain was determined by X-ray crystallography at 3.15 Å resolution, to reveal as expected three β-stranded repeat motifs arranged in a prism (Fig 1A; see Table 1 for data collection and refinement statistics). Each repeat consists of an inner four-stranded β-sheet and an outer two-stranded β-sheet, locked together by two disulfide bonds. Comparison with TgPLP1 APCβ domain (root mean standard deviation, RMSD = 0.978Å between 170 atom pairs) reflects a conserved C-terminal domain architecture across the *Toxoplasma* and *Plasmodium* APCβ domains (Fig S1). Flexible loops between the inner and outer beta-sheets in the third repeat, which are absent in the crystal structure, were subsequently modelled in for molecular dynamics (MD) studies (see Materials and Methods below). Three 100 ns atomistic MD simulations of the PvPLP1 APCβ domain structure indicated that the overall structural core was stable, at an average RMSD of 2.5 Å, while the modelled loops were more flexible (Fig S2).

**Fig 1:**
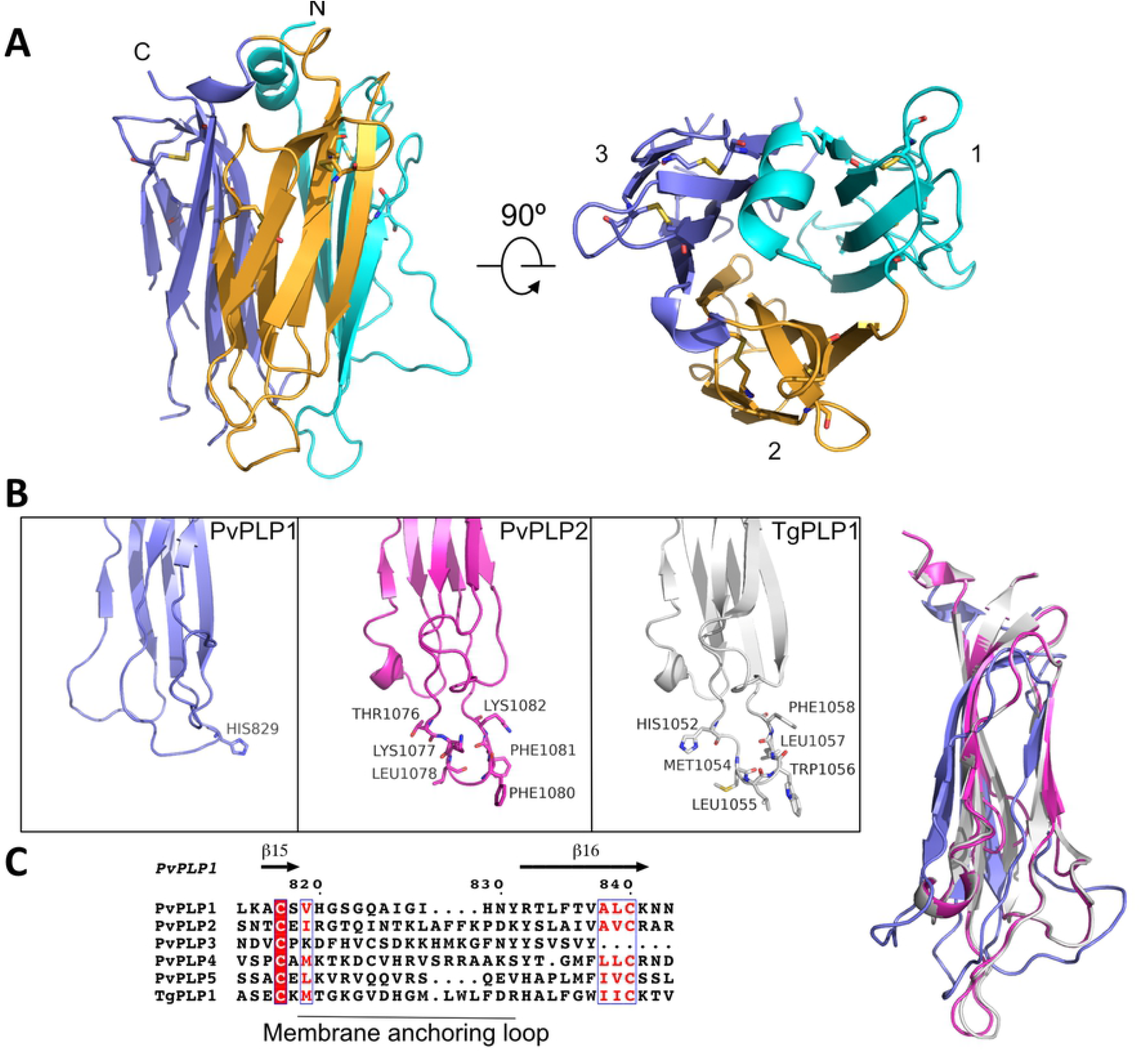
The crystal structure of PvPLP1 APCβ domain and structural comparison with other ApiPLPs. (A) Front and top view of the 3.15Å crystal structure of PvPLP1 APCβ. The structure is composed of three repeats of a β-sandwich fold, each colored in cyan, orange and purple. Density for the turns at the base of repeat 3 was missing. (B) Boxes, comparison of the structures of the PvPLP1 and TgPLP1 APCβ domain with the homology model of the PvPLP2 APCβ, focusing on the third beta-sandwich module and highlighting hydrophobic and basic residues in the proposed membrane anchoring amphipathic loop. On the right is shown a superposition of the three structures, highlighting the agreement between the TgPLP1 and PvPLP2 models. (C) Sequence alignments of *Plasmodium vivax P*PLPs and TgPLP1 amphipathic loop sequences shows low sequence conservation at this region, and truncation of the third repeat in PvPLP3.

**Table 1.** Data collection and refinement statistics.

|  |  |
| --- | --- |
| Wavelength | 0.9686 |
| Resolution range | 63.01 - 3.15 (3.263 - 3.15) |
| Space group | P 1 21 1 |
| Unit cell | 73.74 42.02 87.22 90 104.95 90 |
| Total reflections | 122538 (2207) |
| Unique reflections | 9099 (790) |
| Multiplicity | 13.5 (2.8) |
| Completeness (%) | 98.40 (84.67) |
| Mean I/sigma(I) | 11.32 (1.68) |
| Wilson B-factor | 85.91 |
| R-merge | 0.1942 (0.5668) |
| R-meas | 0.201 (0.6824) |
| R-pim | 0.04997 (0.371) |
| CC1/2 | 0.995 (0.695) |
| CC* | 0.999 (0.906) |
| Reflections used in refinement | 9098 (790) |
| Reflections used for R-free | 431 (37) |
| R-work | 0.2134 (0.3271) |
| R-free | 0.2715 (0.4303) |
| CC(work) | 0.936 (0.742) |
| CC(free) | 0.909 (0.015) |
| Number of non-hydrogen atoms | 3684 |
| Protein residues | 483 |
| RMS(bonds) | 0.012 |
| RMS(angles) | 1.44 |
| Ramachandran favored (%) | 86.12 |
| Ramachandran allowed (%) | 13.23 |
| Ramachandran outliers (%) | 0.65 |
| Rotamer outliers (%) | 0.47 |
| Clashscore | 16.73 |
| Average B-factor | 84.11 |

Statistics for the highest-resolution shell are shown in parentheses.

The loops at the base of TgPLP1 APCβ, especially the extended tryptophan-tipped hydrophobic loop (“dagger”) are important for anchoring the protein to the membrane [30, 31]. The overall sequence identity among *Plasmodium vivax* PLPs is very low, with an average of 20.2 % (Fig S3 and Fig S4). Residues in the membrane binding loop of TgPLP1 APCβ [30, 31], although poorly conserved across all ApiPLPs, are significantly more conserved within each ApiPLP individually, indicating protein-specific functional importance (Fig S4C). Comparison of loops of APCβ in PvPLP1and TgPLP1 reveals that PvPLP1 lacks an extended loop and any aromatic residues at that point in the structure, containing instead several polar and aliphatic hydrophobic residues, as well as a histidine residue at its end rather than a tryptophan. In order to enable a side-by-side comparison of PvPLP1 and PvPLP2 we produced a homology model of PvPLP2 APCβ using the TgPLP1 and PvPLP1 structures, facilitated by conservation of cysteine residues and hydrophobic core residues (Fig S1). The homologous membrane binding loop in PvPLP2 (^1077^*KLAFFK^1082^*) is longer than TgPLP1, but similarly tipped by two aromatic residues (Fig 1B). The hydrophobic and basic residues in the loop may contribute to membrane anchoring as it is amphipathic (later discussed as “loop 1”).

Modelling of PvPLP4 and PvPLP5 was possible on the same basis as PvPLP2, however calculation of sequence identity of each β-sandwich repeat separately found the third repeat to be the most variable, and its truncation in the PvPLP3 APCβ (Fig 1C and Fig S3) meant we could not confidently generate a model of its structure. The PvPLP4 and PvPLP5 APCβ models indicated an absence of the key hydrophobic loop compared to those of PvPLP1 and PvPLP2 (Fig S5B). We nevertheless hypothesise that the function of the APCβ domain is conserved across all PLPs as a membrane binding module, allowing for adaptation to the diverse range of cellular environments within which *Plasmodium* PLPs function, and that their sequence and structural variation in this region are likely to relate to different binding preferences (Fig 1B). Also, the possible formation of a membrane-targeting complex by PvPLPs 3-5 means that they all do not need to have membrane recognition or binding properties, as is also the case with the complement membrane attack complex component proteins [12].

### PvPLP1 does not bind to membranes, while PvPLP2 binds in two distinct orientations

The membrane binding capacity of PvPLP1 and PvPLP2 APCβ domains was probed firstly by lipid dot-blot assays. PvPLP1 did not bind to any mammalian membrane lipid species tested while PvPLP2 recognized anionic lipid species, with a strong preference for phosphatidylinositol 4-phosphate (PI(4)P), followed by palmitoyloleoylphosphatidylserine (PS) and Cardiolipin, and a weak association with the other phosphatidylinositol phosphates (PIPs) (Fig 2). These results were confirmed by liposome sedimentation assays. Affinity of PvPLP1 and 2 for membranes containing Cholesterol, Sphingomyelin (SM), Cardiolipin, PS, L-α-phosphatidic acid (Egg-PA), L-α-lysophosphatidylserine (Lyso-PS) and brain and liver total lipid extract were tested. PvPLP1 showed no binding to any membranes, while PvPLP2 bound to membranes containing negatively-charged lipid species like Cardiolipin, PS and Egg-PA (Fig 2B and C).

**Fig 2:**
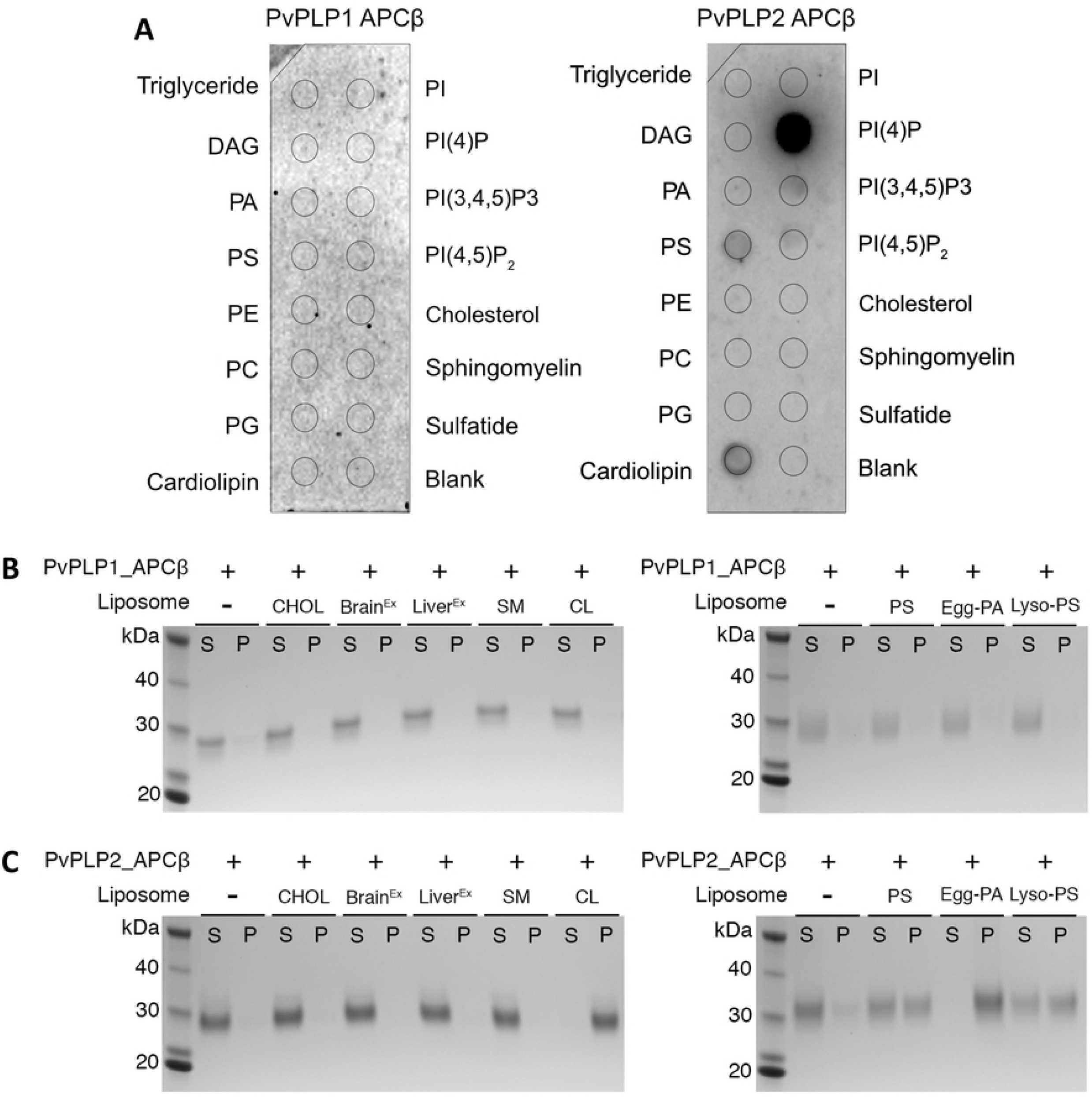
PvPLP1 and PvPLP2 APCβ show different membrane binding capacities. (A) Representative lipid dot-blots. APCβs were incubated with commercially available nitrocellulose membrane dotted with common mammalian membrane lipid species. (B-C) Liposome sedimentation assays of APCβ domains of PvPLP1 (B) and PvPLP2 (C). All liposomes are composed of 40% phosphatidylcholine (PC), 30% phosphatidylethanolamine (PE) and 30% of varied lipids. CHOL, Cholesterol; Brain^Ex^, Brain Total lipid Extract; Liver^Ex^, Liver Total Lipid Extract; SM, Sphingomyelin; CL, cardiolipin; PS, phosphatidylserine; Egg-PA, L-α-phosphatidic acid; Lyso-PS, L-α-lysophosphatidylserine.

Membrane association simulations were next performed to further probe APCβ membrane binding characteristics. PvPLP1 and PvPLP2 APCβs were positioned above a bilayer containing lipids palmitoyloleoylphosphatidylcholine (PC), palmitoyloleoylethanolamine (PE) and PS (45:45:10) representative in simple terms of the eukaryotic membrane inner leaflet, and allowed freely to associate with the membrane over 2 μs in coarse grained (CG) Martini simulations. Sets of 20 simulations for each protein were performed. Only two binding events were observed in PvPLP1 simulations (Fig 3 and Fig S6). In one case, transient binding lasted for 500 ns, where PvPLP1 bound in a tilted orientation facilitated by association with the long loop between β11 and β12 in the third repeat. In the other case, PvPLP1 bound in an upside-down orientation, with the top corner of the domain associated with the bilayer. Given that the MACPF domain would sit above the domain at this location due to the short linker between the two domains, this orientation was deemed non-physiological, as in a previous study with the isolated TgPLP1 APCβ domain [31]. In contrast, PvPLP2 bound to the bilayer in 11 out of 20 (55%) of simulations, in two different orientations, both seemingly of physiological relevance (Fig 3B). In the primary binding orientation (73% of binding events), PvPLP2 sits upright on the bilayer with the phenylalanine-tipped loop at its base anchoring it within the bilayer. This orientation is analogous to that seen in previous studies of TgPLP1 [31]. In the second orientation PvPLP2 lies along the membrane surface with its longitudinal axis parallel to the bilayer (27% of binding events). Visual inspection and lipid contact analysis indicates that for this pose the first tandem repeat of the APCβ domain fold consistently lies along the face of the bilayer, suggesting a specific interaction of this repeat (Fig S6B).

**Fig 3:**
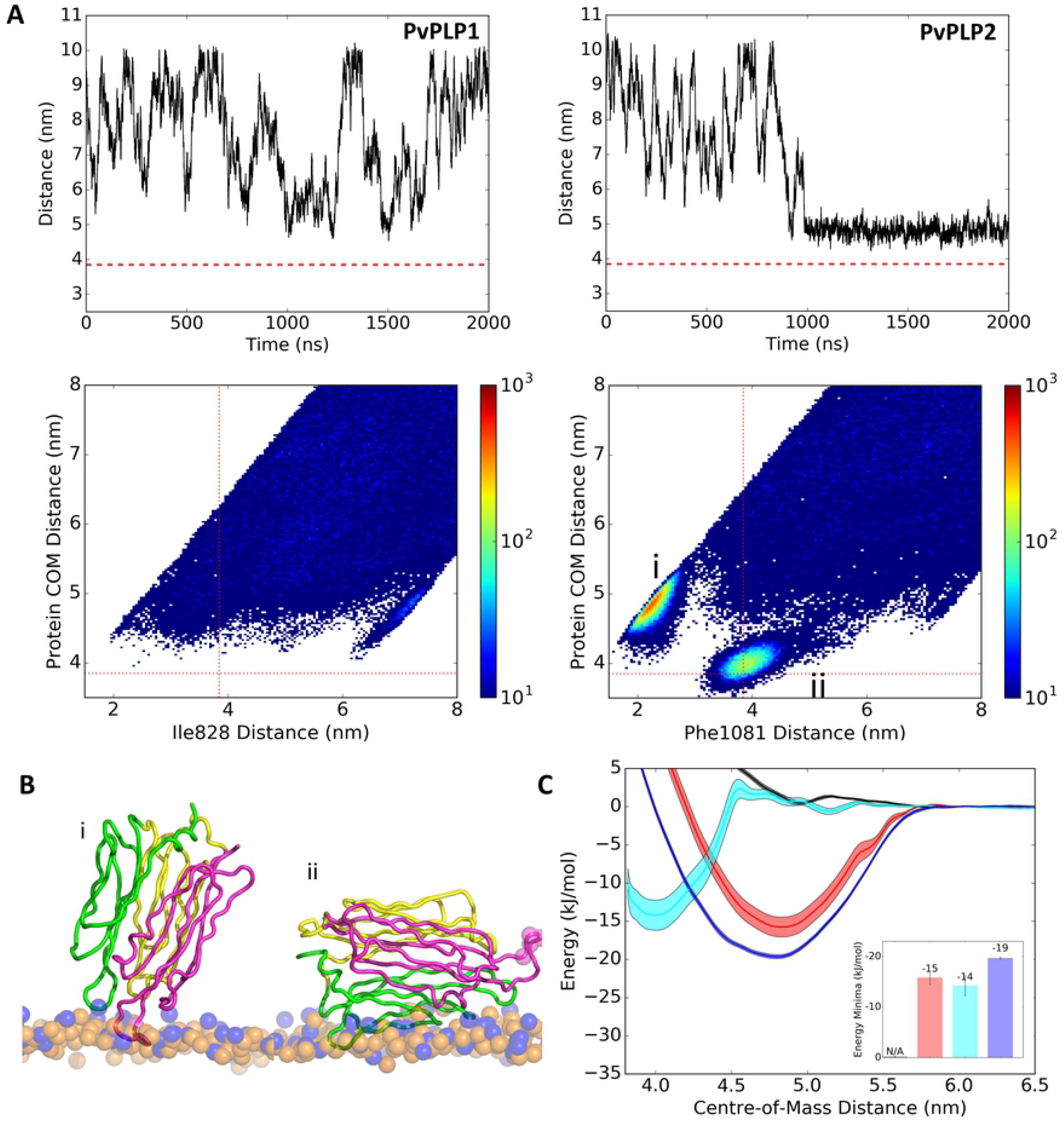
PvPLP1 does not bind to PC:PE:PS (45:45:10) bilayers in simulations while PvPLP2 reveals two distinct binding poses. (A) Top panels: Representative time-distance plots of PLP membrane association simulations. Distance was calculated between the center of mass (COM) of the protein and bilayer. The dashed red line indicates lipid head phosphate distance from COM of membrane. Bottom panels: Density plots indicating COM distance and selected tip residue of the amphipathic loop to bilayer COM distance. Plots show cumulative data of all 20 repeats of each simulation type. (B) Representations of two binding poses identified for PvPLP2 APCβ from simulations. (C) PMF profile for PvPLP1(black), PvPLP2 upright orientation (red), PvPLP2 side orientation (cyan) and TgPLP1 (blue) APCβ domains bound to PC:PE:PS membranes. Distance is measured between COM of the protein and bilayer. Error estimates were obtained using bootstrap analysis. Computed free energy values at the energy minima for each profile are inset.

### Potential of mean force calculation for PvPLPs APCβ binding to membrane

As unbiased MD simulations suggested preference for PvPLP2 binding to the membrane when compared with PvPLP1, we calculated the free energy of membrane binding by PvPLP1 and PvPLP2. For comparison, the same procedure was also performed with TgPLP1. The free energy of binding to PC:PE:PS (45:45:10) bilayers was calculated using orientations taken from the membrane association simulations. Energies were calculated using umbrella sampling simulation methods to measure the potential of mean force (PMF) for binding events. In this approach, each membrane binding domain is placed at a series of positions along a 1-dimensional pathway defining an approach to the membrane long which the PMF will be calculated, and the reaction coordinate path is split into windows each of which is sampled in a separate simulation [33] (Fig 4A). As PvPLP1 did not bind to the membrane in the orientations defined by PvPLP2 and TgPLP1, the protein was aligned with TgPLP1 on the membrane to generate a comparable binding orientation. Similar binding energies of −15 and −19 kJ mol^−1^ were obtained for PvPLP2 and TgPLP1 respectively, while the absence of an energy well for PvPLP1 indicates it has no affinity for the PC:PE:PS (45:45:10) bilayer (Fig 3C). The fact that PMF calculations showed similar free energy of binding of PvPLP2 in a sideways orientation to that of the upright primary binding pose again indicates that they are both of biological relevance, though the greater occupancy of the upright state suggests it is the ultimate and preferred orientation, especially since it is based on a significantly smaller interaction surface area than the sideways orientation.

**Fig 4:**
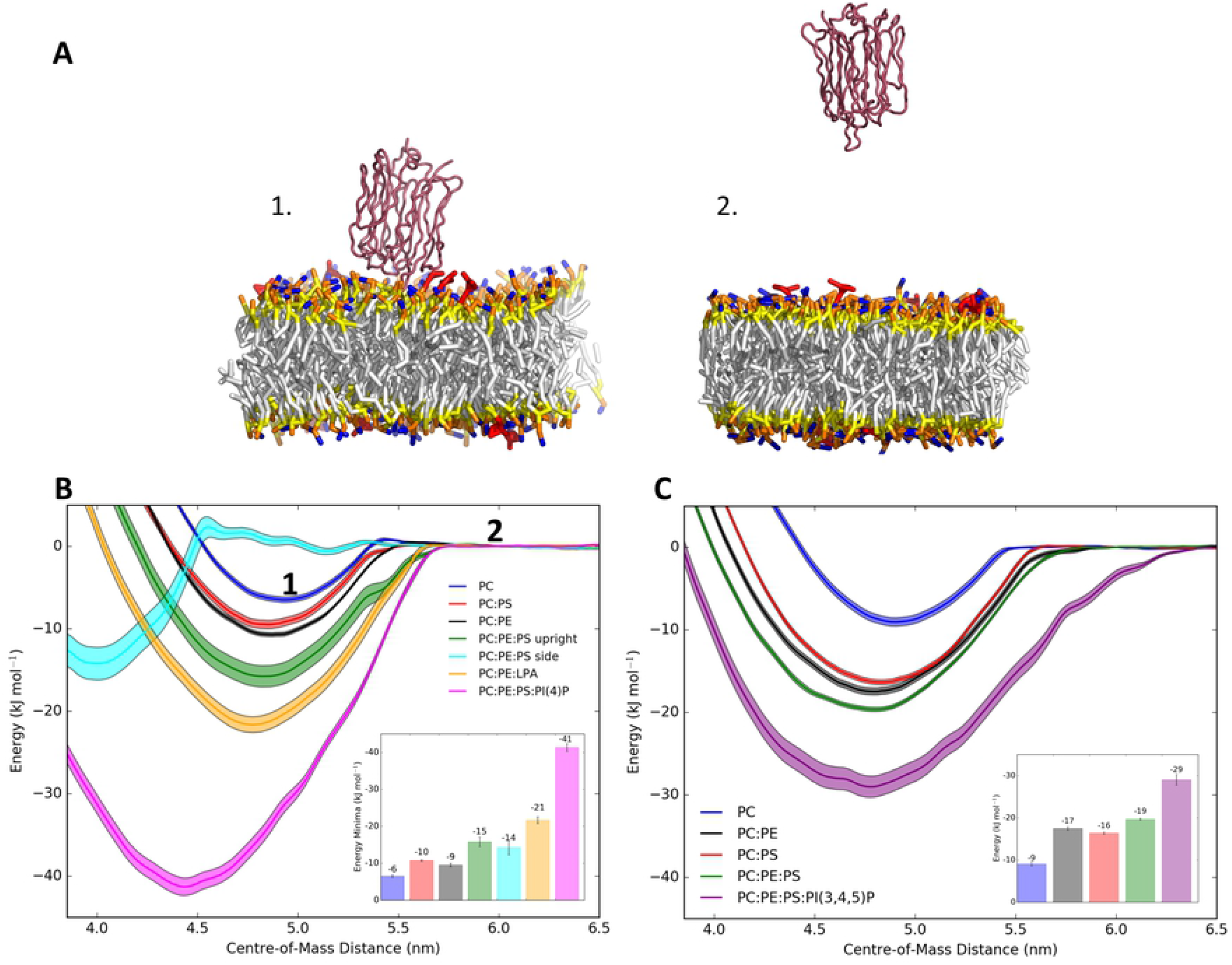
PvPLP2 lipid preferences and mutations within the membrane binding loop. (A) Cartoon representations of PvPLP2 APCβ in the upright orientation bound to the PC:PE:PS (45:45:10) bilayer. Steered MD was used to pull PvPLP2 away from the initial position (1) on the bilayer into solvent (2) to generate umbrella sampling windows. (B) PMF profiles of PvPLP2 APCβ bound to several membranes of different lipid compositions. Positions (1) and (2) are marked. PC:PE and PC:PS were in 80:20 ratios, PC:PE:PS and PC:PE:LPA (Egg-PA) in 45:45:10 ratios, and PC:PE:PS:PI(4)P in the ratio 45:42:10:3. (C) PMF profiles of TgPLP1 APCβ on several bilayers of different lipid composition: PC:PE and PC:PS were 80:20, PC:PE:PS 45:45:10 and PC:PE:PS:PI(3,4,5)P 45:42:10:3.

### Lipid preferences of PvPLP2 by PMF calculations

To explore the lipid preference of PLPs further, we measured free energy of binding on bilayers of different lipid compositions. PMF calculations were first performed with 100% PC, and PC:PE and PC:PS in 80:20 ratio. This was then complemented with two negatively charged lipids, in bilayers of PC:PE:PS (as above) and PC:PE:Egg-PA (molar ratio 45:45:10). PvPLP2 had a low affinity for PC bilayers alone, with addition of PE and PS increasing free energy of binding by −4 and −3 kJ mol^−1^ respectively (Fig 4B). Addition of both PS and PE complement each other to increase binding energy by a further −5 kJ mol^−1^. PvPLP2 had a greater affinity for Egg-PA (LPA) over PS, increasing binding energy by −12 kJ mol^−1^ over PC:PE alone. Accordingly, PC, PE and PS and Egg-PA all contribute to binding of PvPLP2, and complementation with both PE and a negatively charged lipid improved binding affinity. Umbrella sampling simulations with a PC:PE:PS:PI(4)P bilayer (molar ratio 45:42:10:3) were also performed. Free energy of binding of PvPLP2 to a PI(4)P containing bilayer was −41 kJ mol^−1^, a substantial increase in energy of −26 kJ mol^−1^ over the PC:PE:PS bilayer consistent with the lipid dot blot result (Fig 2A).

In comparison, TgPLP1 had a similar increase in binding energy with the addition of either PE and PS of −8 and −7 kJ mol^−1^, respectively (Fig 4C). However, presence of both PS and PE only increased binding energy by ~-3 kJ mol^−1^ for TgPLP1. Our previous work found that TgPLP1 showed a preference for phosphatidylinositol 3,4,5-triphosphate (PI(3,4,5)P_3_) and phosphatidylinositol 4,5-bisphosphate (PI(4,5)P_2_) containing bilayers [31]. Consistently, PMF calculations with a PC:PE:PS:PI(3,4,5)P_3_ bilayer (molar ratio 45:42:10:3) showed an increase in affinity of −10 kJ mol^−1^ over a PC:PE:PS bilayer.

Taken together, these results indicate that PvPLP2 and TgPLP1 have an affinity for negatively charged lipids that generally reside in cytoplasmic leaflets. In contrast, PvPLP1 does not bind to membranes under conditions tested here. The significantly increased affinity of −26 kJ mol^−1^ of PvPLP2 to PI(4)P, compared to −10 kJ mol^−1^ for TgPLP1 on PI(3,4,5)P_3_ containing bilayers suggests that electrostatic interactions may play a more important role in PvPLP2 membrane binding than in TgPLP1 [31].

### Aromatic, hydrophobic and basic residues contribute to membrane binding of PvPLP2

To investigate the protein-lipid interactions of PvPLP2 in greater detail, the two orientations were simulated further in full atomistic simulations. Representative systems were taken from CG simulations and converted to atomistic resolution, following which three repeats of 100-ns duration were produced for each orientation. For each orientation the protein had similar overall stability (Fig S7) and remained bound to the membrane. Over the course of one repeat of the upright orientation, PLP2 began to tilt sideways and transit to the sideways orientation. The other two repeats maintained their initial upright position. Simulations in the sideways orientation remained in a consistent orientation throughout the 100 ns. Again, this perhaps relates to the greater buried surface in this binding orientation.

Lipid contact analysis following atomistic simulations reaffirmed that the phenylalanine tipped loop (loop 1) anchored the protein into the bilayer. In this orientation, a secondary lysine-rich loop (^878^KNK^880^, “loop 2”) further stabilized the membrane interactions (Fig 5A). A series of hydrophobic residues protruding from the surface of repeat 1 of the APCβ domain also had high contact with the bilayer in the sideways orientation, with several lysines and an asparagine residue also contributing to the membrane binding (Fig 5B). We termed a concentrated subset of these residues (^903^FGKK^906^) “loop 3”.

**Fig 5:**
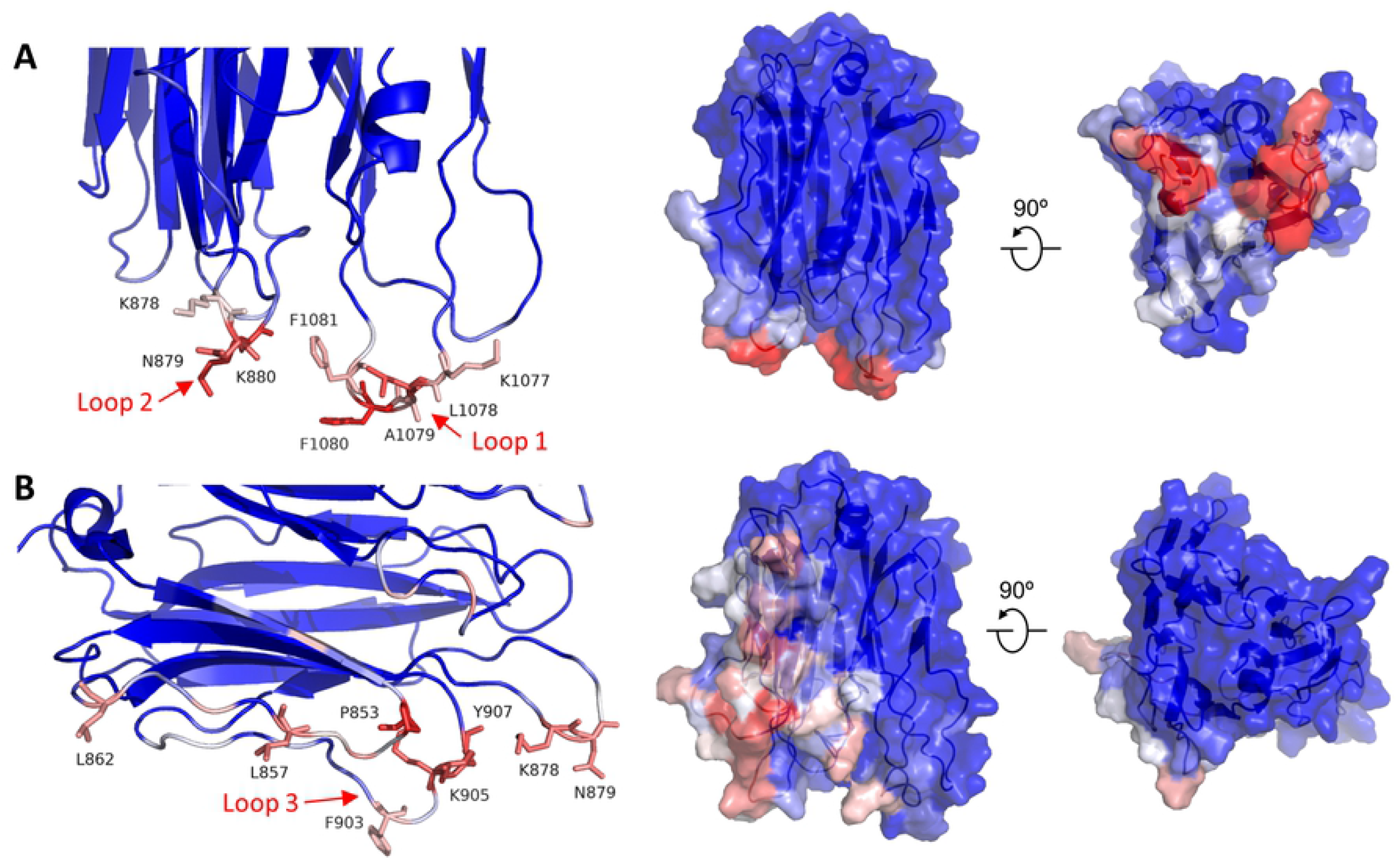
Protein-lipid contacts for 2 orientations of PvPLP2. Cartoon (left) and surface representations (right) of PvPLP2 APCβ, colored by normalised number of lipid contacts averaged over three 100ns atomistic simulations. A: upright orientation, B: side orientation.

To investigate the energetic contribution of these residues to membrane binding, a series of *in silico* mutations were then produced. The equivalent *in silico* mutation of ^1077^KLAFFK^1082^ → AAAAAA (loop 1) entirely abolished membrane affinity in the upright orientation (Fig 6A). This result also occurred when a double mutant of F1080A:F1081A was tested, while mutation of both lysines, K1077 and K1082, in the same loop only minimally reduced binding by 2 kJ mol^−1^ over the wild-type protein. F1080 was identified in lipid contact analysis as the primary phenylalanine responsible for membrane anchoring. Point mutation of this residue to alanine substantially reduced computed membrane affinity to −4 kJ mol^−1^. Atomistic simulations also identified that K880 on loop 2 was frequently in contact with the membrane. K880A showed only a minimal reduction in binding energy. Although the double loop mutant shows a deeper energy well of −17 kJ mol^−1^, this is unrelated to binding to the membrane surface – it does not show the relaxation to 0 kJ mol^−1^ that the wild type protein does and its energetic profile is, like that of K880A and F1080A, truncated. A possible explanation of the energy minimum is the hydrophobic effect as this mutant form of PvPLP2 APCβ replaces 6/7 polar and charged residues with seven alanines (one alanine substitutes a phenylalanine) with consequent effects via water on the free energy of the system. As the tip tryptophan in the extended membrane binding loop of TgPLP1 was identified as a key residue in membrane binding, analogous to F1081 in PvPLP2, a W1056A mutant was compared with PvPLP2 mutants. W1056A reduced the free energy of binding to a PC:PE:PS:PIP3 bilayer by 15 kJ mol^−1^ (Fig 6B), consistent with liposome sedimentation assays which show mutations in this extended loop reduced membrane binding [30, 31].

**Fig 6:**
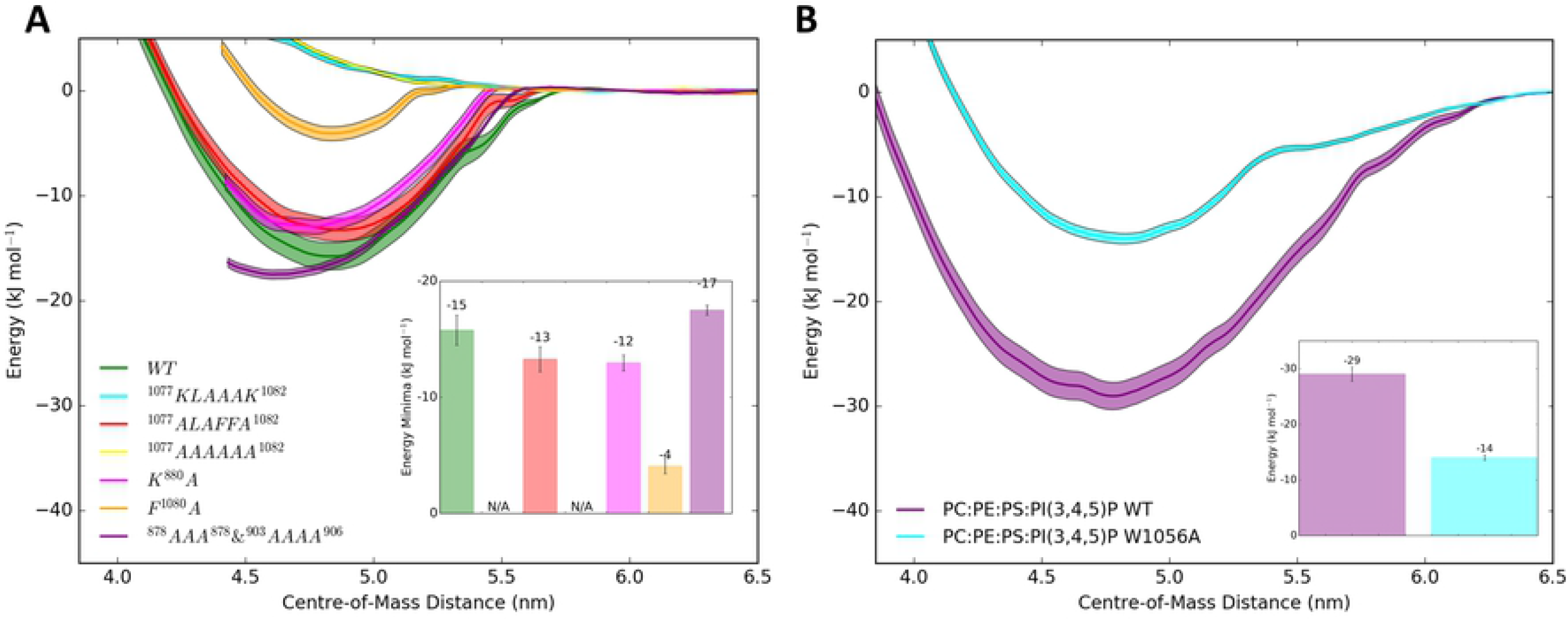
Lipid binding preferences and tryptophan mutation of TgPLP1. (A) PMF profiles for PvPLP2 APCβ wildtype and membrane binding loop mutants FF→AA and KK→AA on bilayers composed of PC, PE and PS (45:45:10). Bootstrap analysis was used to generate error estimates. (B) PMF profiles for TgPLP1 APCβ wildtype and W1056A mutation on a PC:PE:PS:PI(3,4,5)P_3_ (45:42:10:3) bilayer. Bootstrap analysis was used to generate error estimates.

### Mutation of aromatic residues in membrane anchoring loops reduces membrane binding

Following identification, via MD, of the membrane binding role of hydrophobic and positively charged residues at the base of the protein, we produced two full-length PvPLP2 mutants targeting these regions. In the first mutant, the primary membrane binding loop (“loop 1”) was mutated to alanine (^1077^KLAFFK^1082^ → AAAAAA). The second mutant targets the positively charged residues in loop 2 and 3 at the base of PvPLP2, with ^878^KNK^880^ (loop 2) and ^903^FGKK^906^ (loop 3) mutated to alanines. Liposome sedimentation assays with these mutants showed a similar effect: the binding of the PvPLP2 to three different forms of liposomes was substantially reduced compared to that of the wild-type proteins (Fig 7).

**Fig 7:**
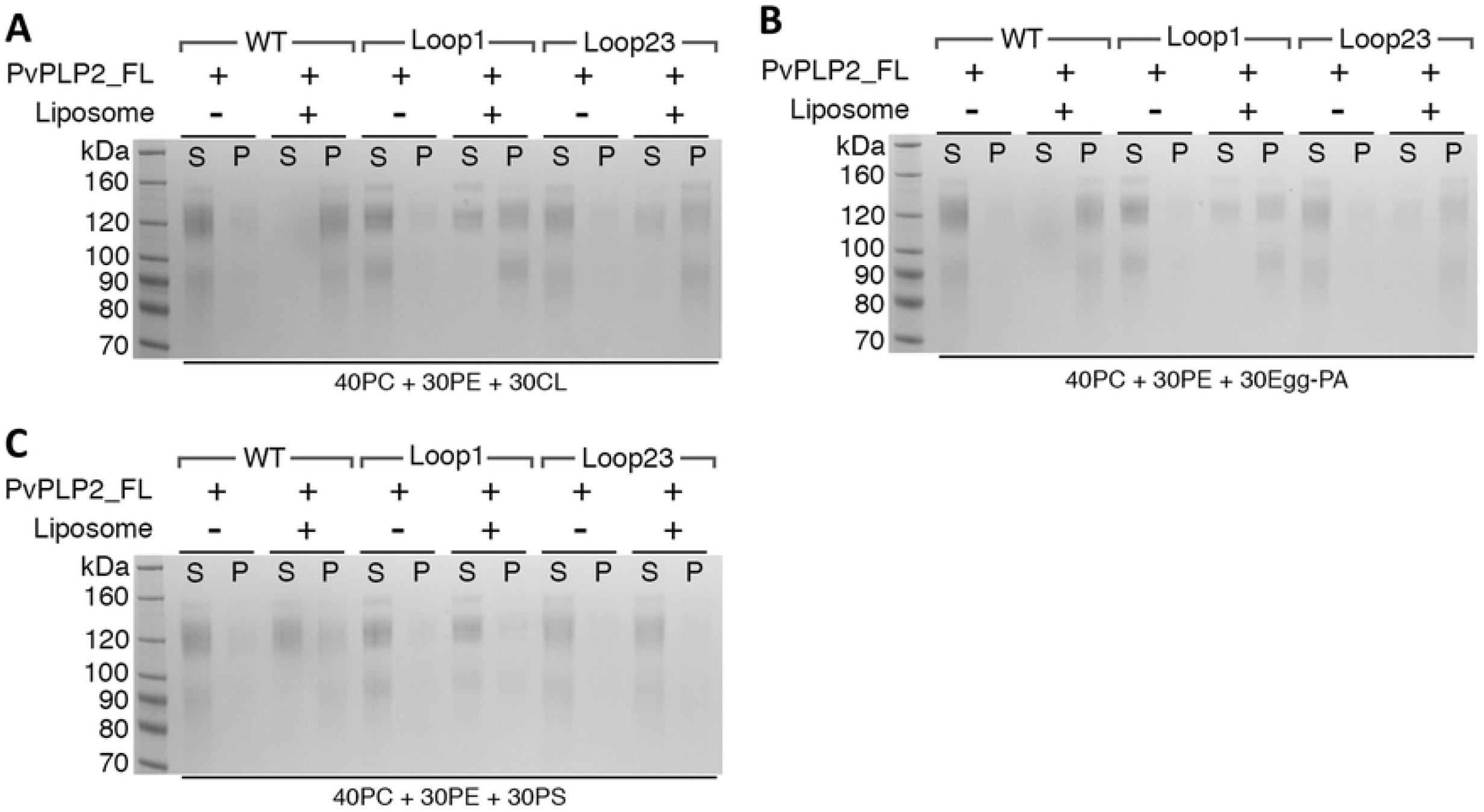
PvPLP2 mutant liposome binding assays. (A-C) Ultracentrifugation-based liposome-binding assays showing that mutations in the hydrophobic tip and charged residues within the APCβ domain (Loop1 and Loop2/3 mutants) reduce the binding of PvPLP2 full-length protein to Cardiolipin (A); L-α-phosphatidic acid (B) and phosphatidylserine (C), indicating the role of APCβ domain for membrane binding of PvPLP2. “Loop1” represents the mutant that affects the Phenylalanine loop tip (^1077^KLAFFK^1082^ - AAAAAA); “Loop2/3” indicates the mutant gives charge neutralization (^878^KNK^880^ - AAA & ^903^FGKK^906^ - AAAA).

## Discussion

PPLPs disrupt membranes during several stages of the *Plasmodium* lifecycle, acting in both cell traversal and egress. Given the diversity of membrane compositions at different stages, PPLPs might have evolved with specific membrane binding characteristics at each stage. Here we show the first crystal structure of a *Plasmodium* PLP C-terminal APCβ domain, that of PvPLP2 which we find to be similar to the TgPLP1 APCβ domain. The similarity in structure, conservation of disulfide locking cysteines and hydrophobic core residues across the PPLP family further confirm a shared domain structure across the ApiPLPs. Combined with MD simulation studies as well as lipid dot blots and liposome sedimentation assays, we show that PvPLP2 APCβ binds to membranes, while PvPLP1 APCβ had little affinity for the membrane lipids. PvPLP1 may rely on a different method for membrane anchoring, possibly via an unidentified glycolipid or a protein receptor, or requiring an unidentified mechanism of activation. This is supported by the distinctly different amino acid composition of the PvPLP1 basal long loop, responsible for primary interaction of PvPLP2 and TgPLP1 with membranes. Similar to PvPLP1, PvPLP4 and 5 lack a phenylalanine/tryptophan containing loop (Fig S5), which indicates that they too may bind to membranes with a different mechanism to PvPLP2 and TgPLP1.

Despite low sequence identity, we showed that PvPLP2 APCβ bound to membranes in a similar mechanism to TgPLP1: an extended loop at the base of the domain tipped with an aromatic residue facilitates primary membrane anchoring for both proteins [31] (Fig 4–7). PvPLP2 had affinity for PE, PS and PI(4)P, which are enriched in the inner leaflet of the plasma membrane. We observed similar affinity towards inner leaflet lipids for TgPLP1, which confirms previous experimental observations [30, 31]. However, PvPLP2 contains additional positively-charged loops (loop 2 and loop 3), which stabilize the primary binding through charge-charge interactions (Fig 7). Consistently, PMF calculations revealed a marked increase in binding energy of PvPLP2 in PI(4)P containing membranes compared to the more moderate increase in TgPLP1 binding upon addition of PI(3,4,5)P_3_ (Fig 4). Mutation of these residues substantially decreased the binding efficiency to negatively-charged membranes (Fig 6 & 7). These findings suggest that anionic lipids may play a more important role in PPLP2 membrane binding, whereas TgPLP1 may have a broader binding profile with less specificity. This could reflect the more confined role of gametocyte egress in which PvPLP2 functions, with PvPLP2 targeting the inner plasma membrane of the cell, compared to the broader replicative niche in which *Toxoplasma* operates.

Interestingly, besides the primary upright membrane anchoring orientation via an extended phenylalanine tipped loop, PvPLP2 also bound to the membrane on its side, an orientation that has not been observed for other PPLPs. PMF calculations showed that PvPLP2 had a similar affinity for the membrane in both orientations, though the more upright state appeared to be the preferred pose. Additionally, loop 2 (^878^KNK^880^) formed a contact point for both orientations and might be able to act as a pivot point between them. Given the preference for the upright pose, despite its smaller buried surface area, we interpret the side-on binding event as a synergy interaction increasing the chance of membrane binding.

This paper provides the first study of the molecular basis of *Plasmodium* PLP APCβ membrane binding. We have revealed structural and functional similarities between members of the ApiPLP family, while also identifying key differences in membrane binding between *Plasmodium* PLP1 and PLP2. PPLPs function in multiple different environments, cell types and in both processes of cell traversal and egress, throughout the *Plasmodium* lifecycle. The differences identified here indicate that membrane binding may be an important regulatory step for PPLPs and varies across the protein family. Regulation of ApiPLP membrane binding is likely to be important for apicomplexans to prevent off-target membrane disruption and damage, especially during motile stages which require microneme secretion without membrane damage, such as cell invasion. Specificity for positively charged inner leaflet lipids, in addition to the role of pH, may prevent binding to the outer surface of the plasma membrane during cell invasion. Extensive further study will be needed to elucidate the different membrane binding functions and mechanisms of action of PPLPs during both cell egress and traversal. As ookinete midgut traversal is a bottleneck in transmission of malaria it will be important to further understand the molecular determinants of function of PvPLPs 3-5.

## Materials and Methods

### Construct cloning and site-directed mutagenesis

In this study, both PvPLP1 (PlasmoDB: PVX_000810) and PvPLP2 sequence (PlasmoDB: PVX_123515) were codon-optimized (by Thermofisher, GeneArt) for a mammalian expression system. The APCβ domains of *PvPLP1* (residues 584 to 843) and *PvPLP2* (residues 836 to 1104) and the full length of *PvPLP2* (residues 25-1104) were cloned into the pHLsec vector [34] using restriction sites (AgeI/KpnI), appending a signal peptide in the N-terminus and a KTHHHHHH tag in the C-terminus of the protein. During protein secretion into the media, the signal peptide was cleaved, leaving an additional three residues Glu-Thr-Gly at the protein N-terminus. Site-directed mutagenesis of the full-length PvPLP2 (loop1 and loop2&3 mutants) was conducted by overlapping polymerase chain reaction (PCR).

### Protein expression and purification

All the PvPLP1 and PvPLP2 proteins were produced recombinantly from mammalian HEK293T cells using a transient transfection and expression protocol as described [30, 31, 35]. To express the APCβ domains of PvPLP1 and 2, the plasmid DNA (12mg) was transfected into 6 liters of HEK293T cells in the presence of the glycosylation inhibitor kifunensine (final concentration of 5 mM) with 12 mg of polyethylenimine. Four to five days post transfection, the cell media containing secreted recombinant protein was harvested by centrifugation at 5000 rpm for 45 minutes to remove the cell debris. The media was filtered through 0.22 μm Amicon filters and dialyzed against a 10× volume of phosphate-buffered saline [10 mM sodium phosphate dibasic, 1.5 mM sodium phosphate monobasic, and 300 mM NaCl, pH 7.5] at 4 °C overnight. The dialyzed media was filtered again and 5 mM imidazole was added to the media before loading onto a Histrap column (GE healthcare). The protein was eluted using a linear imidazole gradient (20 to 500mM) in 20 mM Tris (pH7.5) and 500 mM NaCl over 60 ml. Pooled protein fractions were deglycosylated with endoglycosidase F1 overnight at 4°C and then concentrated for further purification with a size exclusion column (HiLoad 16/60 or 26/60 superdex 75 prep grade columns, GE Healthcare) in a buffer containing 20 mM Hepes (pH 7.5) and 150 mM NaCl. The eluted fraction corresponding to the monomeric species was concentrated to ~ 20 mg/ml for PvPLP1 and ~7 mg/ml for PvPLP2 and used for further studies.

The full-length PvPLP2 proteins (wild type and loop mutants) were expressed and purified in a similar way. In brief, kifunensine was not added during cell transfection and the harvested cell media was concentrated and buffer exchanged with PBS to a final volume of 500 ml as mentioned above. The dialysed medium was loaded to HisTrap HP (GE Healthcare) column and buffered changed with 50 mM MES (pH6.0) and 50 mM NaCl to load onto a cation exchange column (HiTrap™ SP HP, GE Healthcare). A linear salt gradient (50 mM - 1M NaCl) in 50mM MES was used for the elution over 60 ml. The protein fractions were further purified using SEC (Superdex 200) in 20mM Hepes (pH7.5) and 150mM NaCl. Several smaller protein bands appeared in purified PvPLP2 full-length samples, which may be caused by partial proteolysis. The protein was concentrated to ~ 1.5 mg/ml, flash-frozen in liquid nitrogen and stored in −80°C until further use.

### Crystallization, diffraction data collection, and structure determination

Crystallization of PvPLP1 APCβ domain (~20 mg/ml) was set up using a sitting-drop vapor diffusion method in CrystalQuick 96-well plates by mixing 100 nl of protein with 100 nl reservoir solution with equilibration against 95 μl of reservoir. The initial crystals appeared after ~20 days in 20% PEG 3350, 0.2 M Tri-potassium citrate. As the initial crystals were small and highly branched, further optimisation screening using a dilution series of this condition yielded more pin-like crystals. The resulting crystals were harvested with 20% (v/v) glycerol as a cryoprotectant and flash-frozen in liquid nitrogen. The crystal diffraction data were collected on beamline I24 at Diamond Light Source (Didcot, UK). The crystal diffracted to 3.15Å resolution in space group P2_1_. The structure was determined by molecular replacement using TgPLP1 APCβ (PDB 5OUO) as a model, using phenix.phaser [36, 37]. The structure was refined in phenix.refine [38] with manual corrections in Coot [39].

### Lipid dot-blot assay

Commercial membrane lipid strips spotted with 100 pmol of 15 membrane lipids (Echelon Biosciences P-6002) were used for this assay. Membranes were first soaked in PBST (PBS plus 0.1% Tween20), then blocked using 3% (w/v) non-fat dry milk dissolved in PBST for 1 hour at room temperature. The membrane strips were then washed and incubated with 0.5 μg/ml protein in PBST plus 3% milk at 4 °C overnight. Membranes were washed 5 times with PBST for 10 minutes each time and incubated with rabbit anti-penta-Histidine polyclonal antibody for 1 hour. Membranes were washed again as described above, and incubated with secondary anti-rabbit-HRP conjugate for another hour. Following a final washing step of the membrane, the lipid-bound protein was detected by enhanced chemiluminescence (ECL, BIO-RED).

### Liposome sedimentation assay

POPC (1-palmitoyl-2-oleoyl-*sn*-glycero-3-phosphocholine), POPE (1-palmitoyl-2-oleoyl-*sn*-glycero-3-phosphoethanolamine), Cholesterol, Brain Total lipid Extract, Liver Total Lipid Extract, Sphingomyelin (brain, Porcine), Cardiolipin (bovine heart), POPS (1-palmitoyl-2-oleoyl-*sn*-glycero-3-phosphoserine), Egg-PA (L-α-phosphatidic acid, LPA) and Lyso-PS (L-α-lysophosphatidylserine) were purchased from Avanti Polar Lipids Inc. Liposomes with indicated compositions were prepared as described [11]. In brief, different lipids dissolved in chloroform were mixed (1mg in total) and dried in a clean pyrex tube overnight in a desiccator attached to a VARIO-SP diaphragm pump (Vacuubrand). The lipid film was then rehydrated in 0.5 ml solubilisation buffer (20 mM HEPES pH 7.5, 150 mM NaCl) by vigorous vortexing and 5-10 freeze-thaw cycles. The hydrated mixture was extruded through 100 nm polycarbonate membrane (Whatman) for 11 times and the resulting liposomes were stored at 4°C and used within 2 days. To perform the assays, 2.5 μg of protein was incubated with 50 μl liposomes (2 mg/ml) for 1 hour at 37°C. Protein mixed with 50 μl solubilisation buffer was used as a negative control. The protein-buffer/liposome mixtures were then centrifuged at 67,000 rpm in an ultracentrifuge (Optima™ TL with TLA100.4 rotor) for 20 min at 10°C. The supernatant (S) was collected to examine the liposome-unbound proteins. The pellets (P) were resuspended in an equal volume of the supernatant using solubilisation buffer. Then equal volumes of supernatant and pellet were loaded for SDS-PAGE analysis. Each liposome sedimentation assay was performed at least 3 times.

### Bioinformatics

Homologous proteins to PvPLP1 were identified and a multiple sequence alignment produced using HMMER (http://www.hmmer.org/, https://www.ebi.ac.uk/Tools/hmmer/). Conservation scores were mapped onto the structure using the Consurf server [40]. Multiple sequence alignments were generated and plotted using using ESPript 3.0 [41]. Homology models of APCβ domains and missing loop regions from the PvPLP1 APCβ crystal structure were generated using MODELLER software [42].

### Coarse grained MD simulations

Coarse grained (CG) simulations were performed using the Martini forcefield, v2.2 [43, 44] with protein secondary structure predefined used a force constant of 1000 kJ mol^−1^ nm^−2^. The lower elastic bond cut-off used was 0.5 nm, and the upper cut-off was 1.0 nm. Lennard-Jones interactions were shifted to zero between 0.9 and 1.1 nm. The electrostatic potential energy was shifted to zero between 0 and 1.1 nm. To set up CG simulations, the protein atomic structure was first converted into a Martini CG representation [45, 46] using the Martinize script publicly available from Martini [47]. Water and ions were added to solvate the system, at an ion concentration of 150 mM NaCl. The pressure was coupled semi-isotropically using the Berendsen algorithm [48] with a coupling constant of 4ps, and maintained at 1 bar with a compressibility of 5 × 10^−6^ bar^−1^. (Semi-isotropic coupling maintains the pressure separately for xy and z dimensions of the simulations box which is useful for membrane simulations and reduces distortions in the bilayer.) As above, the system was kept at a constant temperature of 310 K coupled using velocity rescaling [49], with protein, water and ions, and lipids coupled separately and a coupling constant of 1ps. The time-step for integration was 20 fs, and particle coordinates were written to the trajectory file every 200 ps.

CG simulations of APCβ domain association with membranes were performed following Yamamoto et al. [50]. Proteins placed 10 nm away from a preformed bilayer were run through 20 simulations with the APCβ domain rotated sequentially around its *x*, *y* and *z* axis to reduce directional bias. Each simulation was run for 1 μs to allow the protein to associate with the membrane. Analysis for each ensemble of association simulations involved production of a 2-dimensional histogram plotting distance between the protein residues and membrane to determine binding orientations. Lipid contacts were also calculated, in which the frequency of contacts between lipids and protein residues were calculated for each simulation and normalised to produce a fingerprint plot of lipid contacts. Contacts were defined as any lipid and residue within 0.6 nm proximity. This distance was reduced to 0.4 nm in the case of lipid contact calculations in atomistic simulations (see below).

### Atomistic MD simulations

Atomistic simulations were performed using Gromacs v5.1 [51, 52] with the Amber 99SB-ILDN force field [53] and the SPC water model [54] with bond lengths and angles constrained using the LINCS algorithm [55]. To reduce computational burden a cut off of 1nm was used for electrostatic and Van der Waal’s interactions; coulombic interactions were also treated using Particle-Mesh Ewald (PME) electrostatic [56]. Constant temperature at 310K was achieved using velocity rescaling, with protein, lipid and solvent (water and ions) separately coupled to an external heat bath [49]. Pressure was maintained at 1 bar with the Parrinello-Rahman barostat with a coupling constant of 1 ps [57]. After initial energy minimization, equilibration was performed with the positions of heavy atoms restrained using a force constant of 1000 kJ mol^−1^ nm^−3^, allowing the solvent to equilibrate around the protein and lipid molecules. Restraints were then removed for final production runs which were produced in triplicate, with each simulation started with random velocities. Time steps of 0.002 ps were used with the writing out of atomic coordinates every 10 ps.

### PMF calculations

Potential of mean force (PMF) calculations driven by umbrella sampling were used to determine the free energy of binding of APCβ domains to membranes [58]. To set up the system, a steered MD simulation was used to pull the protein from a membrane bound conformational state along the membrane normal. A series of overlapping windows along this reaction coordinate were then defined, typically every 0.1 nm, and each window was simulated for 1000 ns. Lipids were fixed along the Z-component of the simulation using a force constant of 10 kJ mol^−1^ nm^−2^. The centre of mass of the protein relative to the membrane was restrained in each simulation with a force constant of 1000 kJ mol^−1^ nm^−2^. The PMF was calculated using GROMACS g_wham [59, 60], with error estimates obtained using bootstrap analysis. All values were shifted so that the PMF value was 0 in bulk water. Convergence was judged based on the change in PMF value calculated across consecutive 100 ns time intervals. All umbrella sampling was performed using Martini CG simulations and produced using GROMACS [61] with protein structure visualisation using the Pymol molecular graphics suite [62]. Data were presented using the Matplotlib Python plotting software [63] and the Seaborn Python data visualization library (https://zenodo.org/record/1313201#.YGCml9zTU2x).

## Data and materials availability

The PvPLP1 APCβ domain crystal structure has been deposited in the RCSB PDB, accession code 7PLN.

## Supporting information captions

**Fig S1: Structural comparison of PvPLP1 and TgPLP1 APCβ crystal structures and PvPLP2 homology model.**

Cartoon representations of each structure are shown with each beta-sandwich repeat coloured differently and labelled from 1-3 in bottom panel. Membrane anchoring loop is highlighted in each structure by as star symbol.

**Fig S2: Structural stability of PvPLP1 crystal structure is stable in MD simulations.**

(A) Alpha helix (blue) and beta-strand (red) secondary structure propensity (top) and RMSF (bottom) and (B) RMSD, averaged from three 100 ns simulations. Errors were calculated as standard deviations. (C) Cartoon putty representation of PvPLP1 crystal structure. Colouring and width of putty represent RMSF calculated for Cα atoms of each residue averaged across the three simulations.

**Fig S3 *Plasmodium vivax* PLP APCβ sequence alignment.** Generated using ESPript 3.0 [41].

**Fig S4 Sequence conservation across PPLPs.**

(A) Conservation of PLP residues mapped onto PvPLP1 APCβ crystal structure with a cartoon and surface representation. Homologous protein to PvPLP1 were identified and a multiple sequence alignment produced using HMMER (http://www.hmmer.org/, https://www.ebi.ac.uk/Tools/hmmer/). Conservation scores were mapped onto the structure using the Consurf server [40].

(B) Sequence identity matrix of PvPLPs and TgPLP1.

(C) Multiple sequence alignment of *Plasmodium* PLP1 (top) and PLP2 (bottom) from key *Plasmodium* species membrane binding loops shows high sequence conservation within each PLP across the species. Generated using ESPript 3.0 [41].

**Fig S5 Structural comparison of PvPLP1, 2, 4, 5 and TgPLP1 APCβ structures.**

(A) Homology models of PvPLP2, 4 and 5 are shown in cartoon representation alongside PvPLP1 and TgPLP1 crystal structures. Each β-sandwich repeat is coloured separately, with the loop homologous to TgPLP1 membrane anchoring loop labelled with a star (⋆). Each pseudo-repeat is labelled in PvPLP1, with all other PLPs following the same pattern.

(B) Close up view of the base of pseudo-repeat 3, highlighting residues predicted to have a role in membrane binding.

**Fig S6 PvPLP1 and PvPLP2 CG membrane association and protein-lipid contacts.**

(A) Plots showing time v. Z distance between center of mass of protein and membrane in each of 20 repeat membrane association simulations on PC:PE:PS (45:45:10) bilayer for PvPLP1 and PvPLP2 APCβ. Each simulation was 2 μs in length, distance was calculated as absolute distance from bilayer to account for periodicity allowing the protein to bind to either side of the bilayer. The red dotted line indicates distance of phosphate lipid head beads from centre of mass of bilayer.

(B) Lipid contact density plot representing normalised lipid contact for each residue from CG membrane association simulations. Each row of the plot represents one simulation out of 20 repeats for each protein.

**Fig S7 Stability of PvPLP2 APCβ from atomistic simulations on PC:PE:PS (45:45:10) bilayer in upright and sideways orientations.**

Representative binding poses for upright and side binding orientations were taken from CG membrane association simulations on PC:PE:PS bilayers, converted into atomistic structure and simulated for 100 ns with three repeats each. Secondary structure propensity is shown for α helices (blue) and β-strands (red) (top). RMSF (middle) and RMSD (bottom) plots are shown with errors calculated as standard deviations. All plots averaged from three repeats.

## References

1. Amino R, Giovannini D, Thiberge S, Gueirard P, Boisson B, Dubremetz JF, et al. Host cell traversal is important for progression of the malaria parasite through the dermis to the liver. Cell host & microbe. 2008;3(2):88–96. PubMed PMID: 18312843.

2. Ishino T, Chinzei Y, Yuda M. A Plasmodium sporozoite protein with a membrane attack complex domain is required for breaching the liver sinusoidal cell layer prior to hepatocyte infection. Cellular microbiology. 2005;7(2):199–208. PubMed PMID: 15659064.

3. Risco-Castillo V, Topcu S, Marinach C, Manzoni G, Bigorgne AE, Briquet S, et al. Malaria Sporozoites Traverse Host Cells within Transient Vacuoles. Cell host & microbe. 2015;18(5):593–603. Epub 2015/11/27. doi: 10.1016/j.chom.2015.10.006. PubMed PMID: 26607162.

4. Kafsack BF, Carruthers VB. Apicomplexan perforin-like proteins. Communicative & integrative biology. 2010;3(1):18–23. Epub 2010/06/12. PubMed PMID: 20539776; PubMed Central PMCID: PMC2881234.

5. Ni T, Gilbert RJC. Repurposing a pore: highly conserved perforin-like proteins with alternative mechanisms. Philosophical transactions of the Royal Society of London Series B, Biological sciences. 2017;372(1726). Epub 2017/06/21. doi: 10.1098/rstb.2016.0212. PubMed PMID: 28630152; PubMed Central PMCID: PMCPMC5483515.

6. Anderluh G, Kisovec M, Krasevec N, Gilbert RJ. Distribution of MACPF/CDC Proteins. Sub-cellular biochemistry. 2014;80:7–30. Epub 2014/05/07. doi: 10.1007/978-94-017-8881-6_2. PubMed PMID: 24798005.

7. Rosado CJ, Kondos S, Bull TE, Kuiper MJ, Law RH, Buckle AM, et al. The MACPF/CDC family of pore-forming toxins. Cellular microbiology. 2008;10(9):1765–74. Epub 2008/06/20. doi: 10.1111/j.1462-5822.2008.01191.x. PubMed PMID: 18564372; PubMed Central PMCID: PMC2654483.

8. Leung C, Hodel AW, Brennan AJ, Lukoyanova N, Tran S, House CM, et al. Real-time visualization of perforin nanopore assembly. Nature nanotechnology. 2017;12(5):467–73. Epub 2017/02/07. doi: 10.1038/nnano.2016.303. PubMed PMID: 28166206.

9. Metkar S, Marchioretto M, Antonini V, Lunelli L, Wang B, Gilbert RJC, et al. Perforin oligomers form arcs in cellular membranes: a locus for intracellular delivery of granzymes. Cell Death Dis. 2015;22:78–85.

10. McCormack RM, de Armas LR, Shiratsuchi M, Fiorentino DG, Olsson ML, Lichtenheld MG, et al. Perforin-2 is essential for intracellular defense of parenchymal cells and phagocytes against pathogenic bacteria. eLife. 2015;4. Epub 2015/09/25. doi: 10.7554/eLife.06508. PubMed PMID: 26402460; PubMed Central PMCID: PMCPMC4626811.

11. Ni T, Jiao F, Yu X, Aden S, Ginger L, Williams SI, et al. Structure and mechanism of bactericidal mammalian perforin-2, an ancient agent of innate immunity. Sci Adv. 2020;6(5):eaax8286. Epub 2020/02/18. doi: 10.1126/sciadv.aax8286. PubMed PMID: 32064340; PubMed Central PMCID: PMCPMC6989145.

12. Serna M, Giles JL, Morgan BP, Bubeck D. Structural basis of complement membrane attack complex formation. Nature communications. 2016;7:10587. Epub 2016/02/05. doi: 10.1038/ncomms10587. PubMed PMID: 26841837; PubMed Central PMCID: PMC4743022.

13. Ellisdon AM, Reboul CF, Panjikar S, Huynh K, Oellig CA, Winter KL, et al. Stonefish toxin defines an ancient branch of the perforin-like superfamily. Proc Natl Acad Sci U S A. 2015;112(50):15360–5. Epub 2015/12/03. doi: 10.1073/pnas.1507622112. PubMed PMID: 26627714; PubMed Central PMCID: PMCPMC4687532.

14. Lukoyanova N, Kondos SC, Farabella I, Law RH, Reboul CF, Caradoc-Davies TT, et al. Conformational Changes during Pore Formation by the Perforin-Related Protein Pleurotolysin. PLoS biology. 2015;13(2):e1002049. Epub 2015/02/06. doi: 10.1371/journal.pbio.1002049. PubMed PMID: 25654333; PubMed Central PMCID: PMC4318580.

15. Thiery J, Lieberman J. Perforin: a key pore-forming protein for immune control of viruses and cancer. Sub-cellular biochemistry. 2014;80:197–220. Epub 2014/05/07. doi: 10.1007/978-94-017-8881-6_10. PubMed PMID: 24798013.

16. Ecker A, Bushell ESC, Tewari R, Sinden RE. Reverse genetics screen identifies six proteins important for malaria development in the mosquito. Mol Microbiol. 2008;70:209–20.

17. Ecker A, Pinto SB, Baker KW, Kafatos FC, Sinden RE. Plasmodium berghei: plasmodium perforin-like protein 5 is required for mosquito midgut invasion in Anopheles stephensi. Exp Parasitol. 2007;116(4):504--8. doi: 10.1016/j.exppara.2007.01.015.

18. Tavares J, Amino R, Menard R. The role of MACPF proteins in the biology of malaria and other apicomplexan parasites. Sub-cellular biochemistry. 2014;80:241–53. Epub 2014/05/07. doi: 10.1007/978-94-017-8881-6_12. PubMed PMID: 24798015.

19. Tavares J, Formaglio P, Thiberge S, Mordelet E, Van Rooijen N, Medvinsky A, et al. Role of host cell traversal by the malaria sporozoite during liver infection. The Journal of experimental medicine. 2013;210(5):905–15. Epub 2013/04/24. doi: 10.1084/jem.20121130. PubMed PMID: 23610126; PubMed Central PMCID: PMC3646492.

20. Yang ASP, O’Neill MT, Jennison C, Lopaticki S, Allison CC, Armistead JS, et al. Cell Traversal Activity Is Important for Plasmodium falciparum Liver Infection in Humanized Mice. Cell Reports. 2017;18(13):3105--16. doi: 10.1016/j.celrep.2017.03.017.

21. Garg S, Agarwal S, Kumar S, Yazdani SS, Chitnis CE, Singh S. Calcium-dependent permeabilization of erythrocytes by a perforin-like protein during egress of malaria parasites. Nature communications. 2013;4:1736. Epub 2013/04/18. doi: 10.1038/ncomms2725. PubMed PMID: 23591903.

22. Deligianni E, Silmon de Monerri NC, McMillan PJ, Bertuccini L, Superti F, Manola M, et al. Essential role of Plasmodium perforin-like protein 4 in ookinete midgut passage. PLOS ONE. 2018;13(8):1–20. doi: 10.1371/journal.pone.0201651.

23. Wirth CC, Glushakova S, Scheuermayer M, Repnik U, Garg S, Schaack D, et al. Perforin-like protein PPLP2 permeabilizes the red blood cell membrane during egress of Plasmodium falciparum gametocytes. Cellular microbiology. 2014;16(5):709--33. doi: 10.1111/cmi.12288.

24. Hall N, Karras M, Raine JD, Carlton JM, Kooij TW, Berriman M, et al. A comprehensive survey of the Plasmodium life cycle by genomic, transcriptomic, and proteomic analyses. Science (New York, NY. 2005;307(5706):82–6. PubMed PMID: 15637271.

25. Kadota K, Ishino T, Matsuyama T, Chinzei Y, Yuda M. Essential role of membrane-attack protein in malarial transmission to mosquito host. Proc Natl Acad Sci U S A. 2004;101(46):16310–5. PubMed PMID: 15520375.

26. Raibaud A, Brahimi K, Roth CW, Brey PT, Faust DM. Differential gene expression in the ookinete stage of the malaria parasite Plasmodium berghei. Molecular and biochemical parasitology. 2006;150(1):107–13. PubMed PMID: 16908078.

27. Kaiser K, Camargo N, Coppens I, Morrisey JM, Vaidya AB, Kappe SHI. A member of a conserved Plasmodium protein family with membrane-attack complex/perforin (MACPF)-like domains localizes to the micronemes of sporozoites. Molecular and biochemical parasitology. 2004;133(1):15--26.

28. Wirth CC, Bennink S, Scheuermayer M, Fischer R, Pradel G. Perforin-like protein PPLP4 is crucial for mosquito midgut infection by Plasmodium falciparum. Molecular and biochemical parasitology. 2015;201(2):90--9. doi: 10.1016/j.molbiopara.2015.06.005.

29. Kafsack BFC, Pena JDO, Coppens I, Ravindran S, Boothroyd JC, Carruthers VB. Rapid membrane disruption by a perforin-like protein facilitates parasite exit from host cells. Science (New York, NY. 2009;323(5913):530--3. doi: 10.1126/science.1165740.

30. Guerra AJ, Zhang O, Bahr CME, Huynh M-H, DelProposto J, Brown WC, et al. Structural basis of Toxoplasma gondii perforin-like protein 1 membrane interaction and activity during egress. PLOS Pathogens. 2018;14(12):1–21. doi: 10.1371/journal.ppat.1007476.

31. Ni T, Williams SI, Rezelj S, Anderluh G, Harlos K, Stansfeld PJ, et al. Structures of monomeric and oligomeric forms of the Toxoplasma gondii perforin-like protein 1. Science Advances. 2018;4(3). doi: 10.1126/sciadv.aaq0762.

32. Gilbert RJC. Structural Features of Cholesterol Dependent Cytolysins and Comparison to other MACPF-domain containing proteins. In: Anderluh G, Gilbert RJC, editors. MACPF/CDC proteins - Agents of Defence, Attack and Invasion. Dordrecht/NL: Springer; 2014.

33. Domanski J, Hedger G, Best RB, Stansfeld PJ, Sansom MSP. Convergence and Sampling in Determining Free Energy Landscapes for Membrane Protein Association. J Phys Chem B. 2017;121(15):3364–75. Epub 2016/11/04. doi: 10.1021/acs.jpcb.6b08445. PubMed PMID: 27807980; PubMed Central PMCID: PMCPMC5402295.

34. Aricescu AR, Assenberg R, Bill RM, Busso D, Chang VT, Davis SJ, et al. Eukaryotic expression: developments for structural proteomics. Acta crystallographica. 2006;62(Pt 10):1114–24. PubMed PMID: 17001089.

35. Ni T, Harlos K, Gilbert RJC. Structure of astrotactin-2: a conserved vertebrate-specific and perforin-like membrane protein involved in neuronal development. Open Biology. 2016;6:160053.

36. Adams PD, Afonine PV, Bunkoczi G, Chen VB, Davis IW, Echols N, et al. PHENIX: a comprehensive Python-based system for macromolecular structure solution. Acta crystallographica. 2010;66(Pt 2):213–21. Epub 2010/02/04. doi: 10.1107/S0907444909052925. PubMed PMID: 20124702; PubMed Central PMCID: PMC2815670.

37. McCoy AJ, Grosse-Kunstleve RW, Adams PD, Winn MD, Storoni LC, Read RJ. Phaser crystallographic software. J Appl Crystallogr. 2007;40(Pt 4):658–74. PubMed PMID: 19461840.

38. Afonine PV, Grosse-Kunstleve RW, Echols N, Headd JJ, Moriarty NW, Mustyakimov M, et al. Towards automated crystallographic structure refinement with phenix.refine. Acta crystallographica. 2012;68(Pt 4):352–67. Epub 2012/04/17. doi: 10.1107/S0907444912001308. PubMed PMID: 22505256; PubMed Central PMCID: PMC3322595.

39. Emsley P, Cowtan K. Coot: model-building tools for molecular graphics. Acta crystallographica. 2004;60(Pt 12 Pt 1):2126–32. PubMed PMID: 15572765.

40. Ashkenazy H, Abadi S, Martz E, Chay O, Mayrose I, Pupko T, et al. ConSurf 2016: an improved methodology to estimate and visualize evolutionary conservation in macromolecules. Nucleic Acids Res. 2016;44(W1):W344–W50. PubMed PMID: WOS:000379786800057.

41. Robert X, Gouet P. Deciphering key features in protein structures with the new ENDscript server. Nucleic Acids Res. 2014;42(W1):W320–W4. PubMed PMID: ISI:000339715000053.

42. Sali A, Blundell TL. Comparative protein modelling by satisfaction of spatial restraints. J Mol Biol. 1993;234(3):779–815. PubMed PMID: 8254673.

43. Marrink SJ, Risselada HJ, Yefimov S, Tieleman DP, de Vries AH. The MARTINI force field: coarse grained model for biomolecular simulations. J Phys Chem B. 2007;111(27):7812–24. doi: 10.1021/jp071097f. PubMed PMID: 17569554.

44. Monticelli L, Kandasamy SK, Periole X, Larson RG, Tieleman DP, Marrink SJ. The MARTINI Coarse-Grained Force Field: Extension to Proteins. J Chem Theory Comput. 2008;4(5):819–34. doi: 10.1021/ct700324x. PubMed PMID: 26621095.

45. Stansfeld PJ, Sansom MS. From Coarse Grained to Atomistic: A Serial Multiscale Approach to Membrane Protein Simulations. J Chem Theory Comput. 2011;7(4):1157–66. doi: 10.1021/ct100569y. PubMed PMID: 26606363.

46. Stansfeld PJ, Sansom MS. Molecular simulation approaches to membrane proteins. Structure. 2011;19(11):1562–72. Epub 2011/11/15. doi: 10.1016/j.str.2011.10.002. PubMed PMID: 22078556.

47. de Jong DH, Singh G, Bennett WFD, Arnarez C, Wassenaar TA, Schafer LV, et al. Improved Parameters for the Martini Coarse-Grained Protein Force Field. Journal of Chemical Theory and Computation. 2013;9(1):687–97. PubMed PMID: WOS:000313378700069.

48. Berendsen HJC, Postma JPM, Vangunsteren WF, Dinola A, Haak JR. Molecular-Dynamics with Coupling to an External Bath. Journal of Chemical Physics. 1984;81(8):3684–90. PubMed PMID: WOS:A1984TQ73500045.

49. Bussi G, Davide D, Parinello M. Canonical Sampling through Velocity Rescaling. J Chem Phys. 2007;126:14101.

50. Yamamoto E, Kalli AC, Yasuoka K, Sansom MSP. Interactions of Pleckstrin Homology Domains with Membranes: Adding Back the Bilayer via High-Throughput Molecular Dynamics. Structure. 2016;24(8):1421–31. doi: 10.1016/j.str.2016.06.002. PubMed PMID: 27427480; PubMed Central PMCID: PMCPMC4975593.

51. Abraham MJ, Murtola T, Schulz R, Pálla S, Smith JC, Hess B, et al. GROMACS: High performance molecular simulations through multi-level parallelism from laptops to supercomputers. SoftwareX. 2015;1-2:19–25.

52. van der Spoel D, Lindahl E, Hess B, Groenhof G, Mark AE, Berendsen HJ. GROMACS: fast, flexible, and free. J Comput Chem. 2005;26(16):1701–18. doi: 10.1002/jcc.20291. PubMed PMID: 16211538.

53. Lindorff-Larsen K, Piana S, Palmo K, Maragakis P, Klepeis JL, Dror RO, et al. Improved side-chain torsion potentials for the Amber ff99SB protein force field. Proteins. 2010;78(8):1950–8. doi: 10.1002/prot.22711. PubMed PMID: 20408171; PubMed Central PMCID: PMCPMC2970904.

54. Berweger CD, Vangunsteren WF, Mullerplathe F. Force-Field Parametrization by Weak-Coupling - Reengineering Spc Water. Chem Phys Lett. 1995;232(5-6):429–36. PubMed PMID: WOS:A1995QE86700003.

55. Hess B, Becker H, Berenedsen HJC, Fraaije JGEM. LINCS: a linear constraint solver for molecular simulations. J Comput Chem. 1997;18:1463–72.

56. Essmann U, Perera L, Berkowitz ML, Darden T, Lee H, Pedersen LG. A Smooth Particle Mesh Ewald Method. J Chem Phys. 1995;103:8577–93.

57. Parrinello M, Rahman A. Polymorphic transitions in single crystals: A new molecular dynamics method J Appl Phys. 1981;52:7182–90.

58. Kästner J. Umbrella Sampling. Wiley Interdisciplinary Reviews: Computational Molecular Science. 2011;1(6):932–42.

59. Hub JS, de Groot BL, van der Spoel D. g_wham-A Free Weighted Histogram Analysis Implementation Including Robust Error and Autocorrelation Estimates. Journal of Chemical Theory and Computation. 2010;6(12):3713–20. PubMed PMID: WOS:000285217000010.

60. Kumar S, Bouzida D, Swendsen RH, Kollman PA, Rosenberg JM. The Weighted Histogram Analysis Method for Free-Energy Calculations on Biomolecules .1. The Method. Journal of Computational Chemistry. 1992;13(8):1011–21. PubMed PMID: WOS:A1992JL80700011.

61. Berendsen HJC, Vanderspoel D, Vandrunen R. Gromacs - a Message-Passing Parallel Molecular-Dynamics Implementation. Comput Phys Commun. 1995;91(1-3):43–56. PubMed PMID: WOS:A1995TF32200004.

62. Ltd S. The PyMOL Molecular Graphics System, Version 1.8. 2015.

63. Hunter JD. Matplotlib: A 2D graphics environment. Comput Sci Eng. 2007;9(3):90–5. PubMed PMID: WOS:000245668100019.

